# Temporal Analysis of Pituitary Transcriptional Dynamics in Mice Models of Hypopituitarism During Postnatal Development

**DOI:** 10.1101/2024.02.26.582133

**Authors:** Juliana Moreira Marques, Claudia Veiga Chang, Nicholas Silvestre Souza Trigueiro, Ricardo Vieira Araujo, Cinthya dos Santos Cerqueira, Lilian Cristina Russo, Bruna Viscardi Azevedo, Berenice Bilharinho de Mendonca, Nicolas Hoch, Luciani Renata Silveira de Carvalho

## Abstract

Congenital hypopituitarism is characterized by deficient pituitary hormone production, affecting growth and development. The molecular mechanisms underlying pituitary development and dysfunction in hypopituitarism remain incompletely understood. We investigated the expression of key pituitary development markers in three mouse models of congenital hypopituitarism, with molecular alterations in the *Prop1, Pou1f1*, and _α_*GSU* genes across critical postnatal developmental stages: neonatal (P0), early postnatal (P7), pubertal (4 weeks), and adult (8 weeks). We assessed mRNA and protein levels of the pituitary stem cell markers (SOX2), proliferation marker (Ki67) and pituitary hormones, correlating these with pituitary function and disease. *Prop1* deficiency led to significant upregulation of *Sox2* and *Hesx1* during early postnatal development and in adulthood, diverging from the relatively stable expression patterns observed in *Pou1f1* and _α_*GSU* mutants. Despite some variations, overall *Sox2* and *Ki67* expression profiles were similar between *Prop1* and *Pou1f1* mutants. *Prop1* mutants exhibited altered pituitary morphology, with increased SOX2-positive cells suggesting disrupted stem cell migration. During the pubertal period, a subset of hormone-producing cells in *Prop1* mutants co-expressed SOX2, indicating differentiation without restoring normal pituitary function. Hormone analysis revealed transient gonadotropin production and secretion during sexual maturation in *Prop1* mutants, without recovery of the hypogonadal phenotype. Our study elucidates the complex transcriptional dynamics of pituitary development markers in mouse models of congenital hypopituitarism, highlighting the pivotal role of *Prop1* in regulating stem cell marker expression. The distinct transcriptional responses in *Prop1* mutants during key developmental windows shed light on the mechanisms of pituitary dysgenesis and the persistent inability to fully recover pituitary function, despite transient hormonal changes during puberty. These insights contribute to a better understanding of pituitary development and dysfunction in congenital hypopituitarism.

## Introduction

The adenohypophysis produces and secretes six hormones in response to hypothalamic stimuli: growth hormone (GH), luteinizing hormone (LH), follicle-stimulating hormone (FSH), thyroid-stimulating hormone (TSH), prolactin (PRL), and adrenocorticotropic hormone (ACTH)(1). These hormones are produced by specific cell types differentiated from pituitary stem cells in response to the activation of several signaling and transcription factors (2). Pituitary stem cells are important for both development and pituitary postnatal plasticity. SRY-box transcription factor 2 (SOX2) is the main pituitary stem/progenitor cell marker (3-7). SOX2-positive pituitary stem cells are located both in the marginal zone remaining from Rathke’s pouch and in clusters within the adult gland parenchyma. They can differentiate into new hormone producing cells in response to demand (8, 9). SOX9 is expressed in adult pituitary stem cells (10, 11), and its expression increases along with the calcium-binding protein S100β (3).

Prophet of Pit1 (PROP1) is a pituitary-specific transcription factor essential for adenohypophysis development and function. It acts as a repressor to suppress Homeobox gene expression in embryonic stem cell 1 (HESX1) activity and as an activator to initiate expression of POU class 1 homeobox 1 (POU1F1, also known as PIT-1) (12-14). PROP1 is expressed in early stages of pituitary development and plays a role in cellular and molecular processes of hormone-producing cells during gland differentiation (15-19). Bi-allelic loss of function variants in *PROP1* are the main cause of congenital pituitary hormonal deficiencies (hypopituitarism), leading to GH, TSH, PRL, and LH/FSH deficiency and, in some cases, ACTH deficiency accounting for up to 50% of familial cases of congenital hypopituitarism (20-22). *POU1F1* is also necessary f or the differentiation of pituitary hormone-producing cell subtypes. It is responsible for differentiating somatotrophs, lactotrophs, and thyrotropes in the anterior pituitary and plays an important role in pituitary growth (2, 12, 19, 23).

Mouse models are essential tools for understanding pituitary development and the role of transcription factors in pituitary cell specification. One of the most important and widely studied models of congenital hypopituitarism is the Ames dwarf mouse, which has a spontaneous, autosomal recessive, pathogenic variant (p.S83P) in PROP1 that abrogates DNA binding (13, 24, 25). Another model of hypopituitarism is the Snell dwarf, a mouse with a spontaneous pathogenic variant (p.W251C) in the POU1F1 transcription factor, which abolishes DNA binding (26). Genetically engineered models are also of great value to study pituitary disease. Mice homozygous for a genetically engineered null allele of *Cga* have congenital hypopituitarism with primary deficiencies in the glycoprotein hormones, TSH, LH and FSH, and secondary GH and PRL deficiencies due to the lack of thyroid hormone (27).

Most studies of pituitary development have focused on the early stages of organogenesis during embryonic mouse development. However, there is substantial pituitary growth and changes in the relative abundance of differentiated cell types in the postnatal period of development. Little attention has been paid to changes that occur during puberty and sexual maturation. In this study, we aimed to characterize the dynamics of stem cell and proliferation marker gene expression in these three mouse models of congenital hypopituitarism.

## Materials and Methods

### Ethical considerations

This study was approved by the Ethics Committee in Research of the University of São Paulo (number 0217/06).

### Mice

The Ames dwarf mice, DF/B-*Prop1*^*df/df*,^ are homozygous for the spontaneous mutation, p.S83P (c.475C>T; ENSMUST00000051159). Mice were originally obtained from Dr. Andrez Bartke and maintained as a non-inbred, mixed stock at the University of Michigan, and donated by Professor Sally Camper to the University of São Paulo. They were genotyped by polymerase chain reaction (PCR) using the primers 5′-GAGCTGGGGAGACCTAAGCTTTGCC-3′ and 5′-GCCCAGATGTCAGGATACTG-3′, followed by PCR product digestion with Van91I (28). For the experiments examining the period of sexual maturation (P30, P40 and P60), mice were weighed, the naso-anal length was measured, mice euthanized using Isofluorane (Isoforine®, Cristália, São Paulo, Brazil). and whole blood, pituitary, and gonads were collected.

Snell dwarf, *Pou1f1*^*dw/dw*^, were obtained from the Jackson Laboratory mice (DW/J-*Mlph*^*ln*^, *Pou1f1*^*dw*^, Jackson laboratory stock number 000643). They are homozygous for the spontaneous mutation, p.W251C (c.900G>T, ENSMUST00000176330), which abrogates the DNA binding and transactivation properties. Mice were donated by Professor Sally Camper to the University of São Paulo and genotyped by Sanger sequencing using the primers 5′-GCTGCTAAGGATGCTCTGG-3′, 5′-CCTTGGAAATAGAGAACAGGC-3′, 5′-ACACTTCGGGGACCCCAGCA-3′, and 5′-CTCTGCCTTCGGTTGCAGGA-3′.

*Cga* knockout mice, with a neomycin resistance gene in exon 3, were generated by homologous recombination in the embryonic stem cells and blastocyst injection at the University of Michigan (31). Chimeras were mated with C57BL/6 mice to produce a line with a mixed genetic background, officially known as B6:129S2-*Cga*^*tm1Sac*^. Mice were donated by Professor Sally Camper to the University of São Paulo (Jackson Laboratory, stock number 002494) and genotyped using the original protocol.

### RNA extraction

Pituitary samples were collected, fresh-frozen in liquid nitrogen, and subsequently stored at –80°C until further analysis. RNA was extracted from pituitary sample pools from wild-type and mutant mice. Six pituitaries (3M:3F) were pooled for samples from newborn mice, postnatal day 0 (P0), and at the end of the first wave of pituitary development at postnatal day 7 (P7). Four pituitaries (2M:2F) were pooled for samples from 4-week-old (4 W) and 8-week-old (8 W) mice. Five individual male pituitaries were used for the sexual maturation experiments (P30, P40 and P60). RNA extraction was performed using the RNeasy Mini Kit (Qiagen, Valencia, CA, USA), according to the manufacturer’s instructions. RNA quality and purity were assessed by two absorbance ratios: 260/230 between 1.8–2.0 and 260/280 > 1.8 were considered acceptable. RNA integrity was determined by examining 28S and 18S RNA degradation in a 1% agarose gel. RNA samples were maintained at –80ºC until further analysis. Three micrograms of RNA were reverse transcribed into cDNA using the High-Capacity cDNA Reverse Transcription kit (Applied Biosystems, Foster City, CA, USA), according to the manufacturer’s protocol.

### Quantitative PCR (qPCR)

Gene expression was determined using the RT^2^ Profiler PCR Array (SYBR ® Green PCR Mastermix, CAPM11273; SABioscences-QIAGEN, Valencia, CA, USA) and SYBR Green qPCR Master Mix (Thermo Fisher Scientific, Waltham, MA, USA), according to the manufacturer’s recommendations. Amplifications were performed on an ABI PRISM 7000 thermal cycler (Applied Biosystems) using the following cycling conditions: 95 °C for 10 min, followed by 40 cycles of 95 °C for 15 s and 60 °C for 1 min. Gene expression was normalized to beta-actin (*Actb*). In the isolated wild-type and mutant data, relative quantification was performed using the ΔCT method, a modified Livak method for target/reference gene correlation (29). The results were expressed by log_2_ ^Δ^^CT^ with mutant relative to wild-type expression. Each analysis was performed in three different pools at least three times.

### Immunofluorescence

Samples were fixed for 24h (P0 and P7) and 2h (4W and 8W) on paraformaldehyde 4% and DMSO 1%. All samples were washed in PBS, dehydrated in a graded series of ethanol. Pituitaries were re-hydrated through a series of graded ethanol washes and incubated in collagenase (1mg/mL) at room temperature for nine minutes and incubated overnight at 4ºC in a blocking buffer. To visualize SOX2, Ki67 and the pituitary hormones GH, TSH and LH, samples were incubated with antibodies against SOX2 (1:1600; ab97959, Abcam, UK), Ki67 (1:250, Dako), GH (1:250; NHPP), beta-TSH (1:250; NHPP) and beta-LH (1:250; NHPP) overnight at 4ºC. Whole mount tissues were counterstained with propidium iodide (1:1000) and the secondary antibody Alexa Fluor®488 (1:1000; ab97959, Abcam, UK) overnight at 4ºC in a dark chamber.

### Quantification of immunofluorescence

All images were captured with LSM 780-NLO (Carl Zeiss, USA) and processed with ZEN Black Microscope and Imaging software (Carl Zeiss, USA). Images were analyzed using FIJI software (version 2.0.0) by selecting a region of interest (ROI) in each image and measuring the area, intensity, and mean gray value, and calculating the corrected total cell fluorescence (CTCF). Each image was normalized using three background ROIs from the same file.

Cells immunostained for SOX2 and hormones were counted after randomly selecting an area in the parenchyma and marginal zone. The number of cells is reported per area, and at least three RO1 were analyzed per sample per marker. Mean values are reported.

### Hormonal serum levels

Blood samples were collected from five mutant and five wild type animals at each period. Serum levels of the pituitary hormones GH, FSH, and LH were measured using the Mouse Pituitary Magnetic Bead Panel (MPTMAG-49K, Merck, USA) according to manufacturer instructions. Fluorescence intensities were determined using Luminex 200TM (Luminex, USA) and processed and analyzed by Milliplex Analyst 3.5.5.0 (Merck, USA) using standard curves and controls.

### Statistical analysis

Statistical analyses were performed in R. Normality and homoscedasticity were analyzed using the Shapiro–Wilk and Levene tests, respectively. When the data achieved these parameters, one-way or two-way analysis of variance (ANOVA) and the post hoc Tukey test were used; when data were not parametric, robust ANOVA and post hoc Dunn tests were performed. Statistical significance was set at p < 0.05.

## Results

### *Prop1* deficiency changes the transcriptional dynamics of pituitary transcription factor

We assessed the expression of genes critical for anterior pituitary development and cell fate commitment during key postnatal time points: at birth (P0), the end of the first wave of pituitary development (P7), during sexual maturation (4W), and in adulthood (8W) (Figure 1, S1; S2) in three different mouse models of congenital hypopituitarism. We observed that *Prop1* deficiency had a pronounced effect on expression of the stem cell marker *Sox2* and *Hesx1* during post-natal development. *Prop1* mutants had significant increases in expression of both genes at P7 and 8W relative to WT and modest decreases in expression at 4W (Figure 1A and B). Expression of the cell proliferation marker *Ki67* (*mKi67*) was not significantly changed in *Prop1* deficient mice, except for P7, when transcripts were reduced (Figure 1C). Generally, *Pou1f1* and αGSU deficient mice exhibit similar levels of *Sox2, Hesx1*, and *Ki67* transcription relative to WT at most ages examined (Figure 1D-I). *Sox2* and *Hesx1* transcripts are elevated at 8W in the *Pou1f1* deficient mice, and *Hesx1* transcripts are reduced at 4W in αGSU deficient mice. Taken together, the most striking effects were observed in *Prop1* deficient mice, which fail to suppress expression of *Sox2* and *Hesx1* (Figure 1J-L).

**Figure 1.**
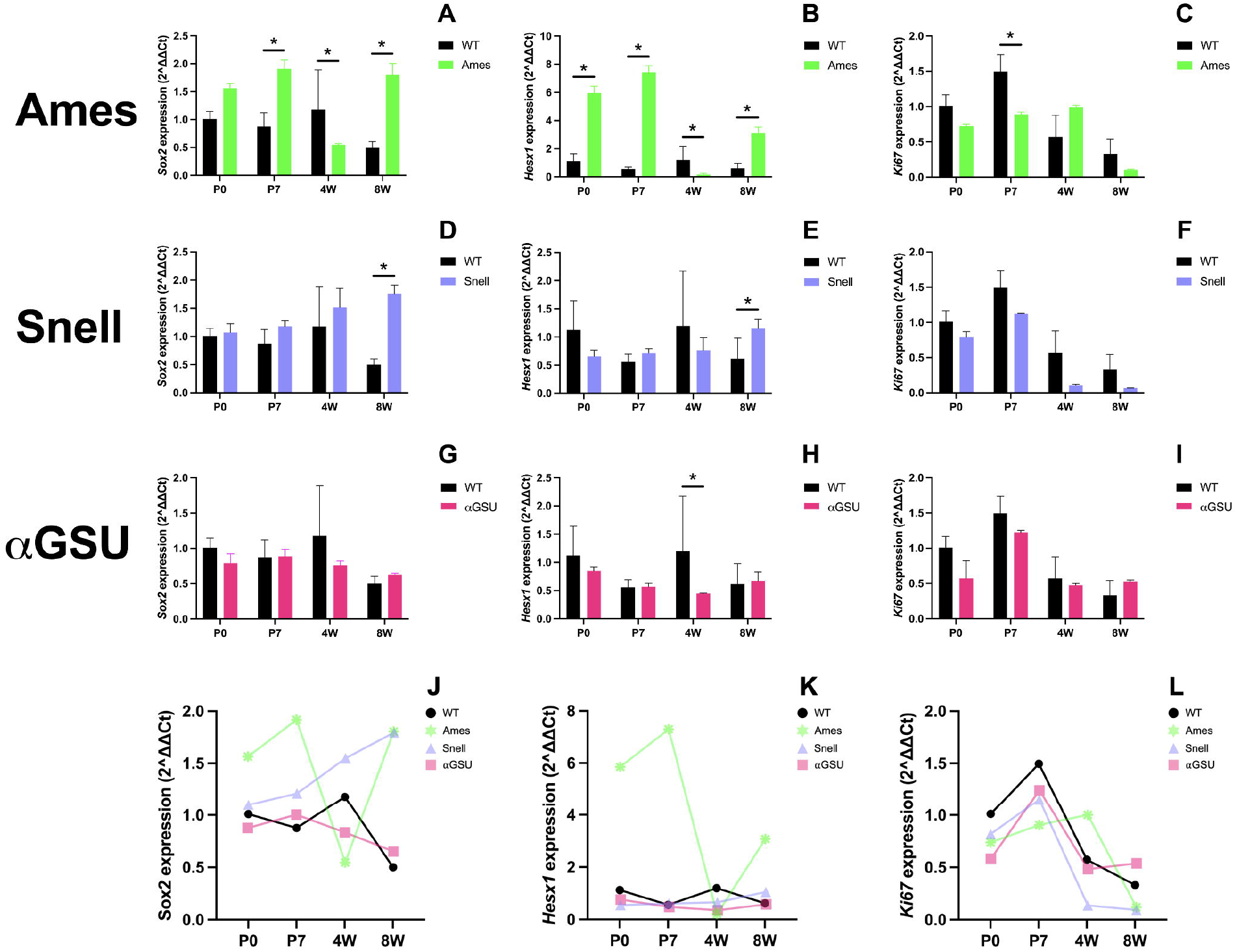
*Sox2* and *Hesx1* transcripts in *Prop1* mutants after birth. RT-qPCR results expressed in log_2_ ^Δ^^CT^ of the genes *Sox2* (A, D and G), *Hesx1* (B, E and H) and *Ki67* (C, F and I) in the wild-type, *Prop1*^*df/df*^, *Pou1f1*^*dw/dw*^, and alpha-glycoprotein subunit (αGSU) mice at postnatal periods (newborn: P0, one week: P7, four weeks: 4W, and eight weeks: 8W). Results are expressed as the mean ± standard deviation (SD). Asterisks indicate significant differences between groups (p < 0.05). Timeline of *Sox2, Hesx1* and *Ki67* expression in the wild-type (black), Ames (green), Snell (purple), and alpha-glycoprotein subunit (αGSU) (pink) mice (J to L) at postnatal periods. Results are expressed as the mean of each group.

### SOX2 expression decreases in the sexual maturation period in *Prop1* and *Pou1f1* mutants

The transcriptional elevation of *Sox2* was prominent in *Prop1* deficient mice at P0, P7, and 8W. Therefore, we examined gene expression at the protein level using immunohistochemical staining for SOX2 and Ki67 (Figure 2). We confirmed that SOX2 protein levels were elevated in *Prop1* mutants at P0, P7 and 8W, and reduced at 4W (Figure 2A). Although the levels of *Ki67* transcripts were reduced at P7 in *Prop1* deficient mice, protein levels were only significantly reduced at 4W (Figure 2B).

**Figure 2.**
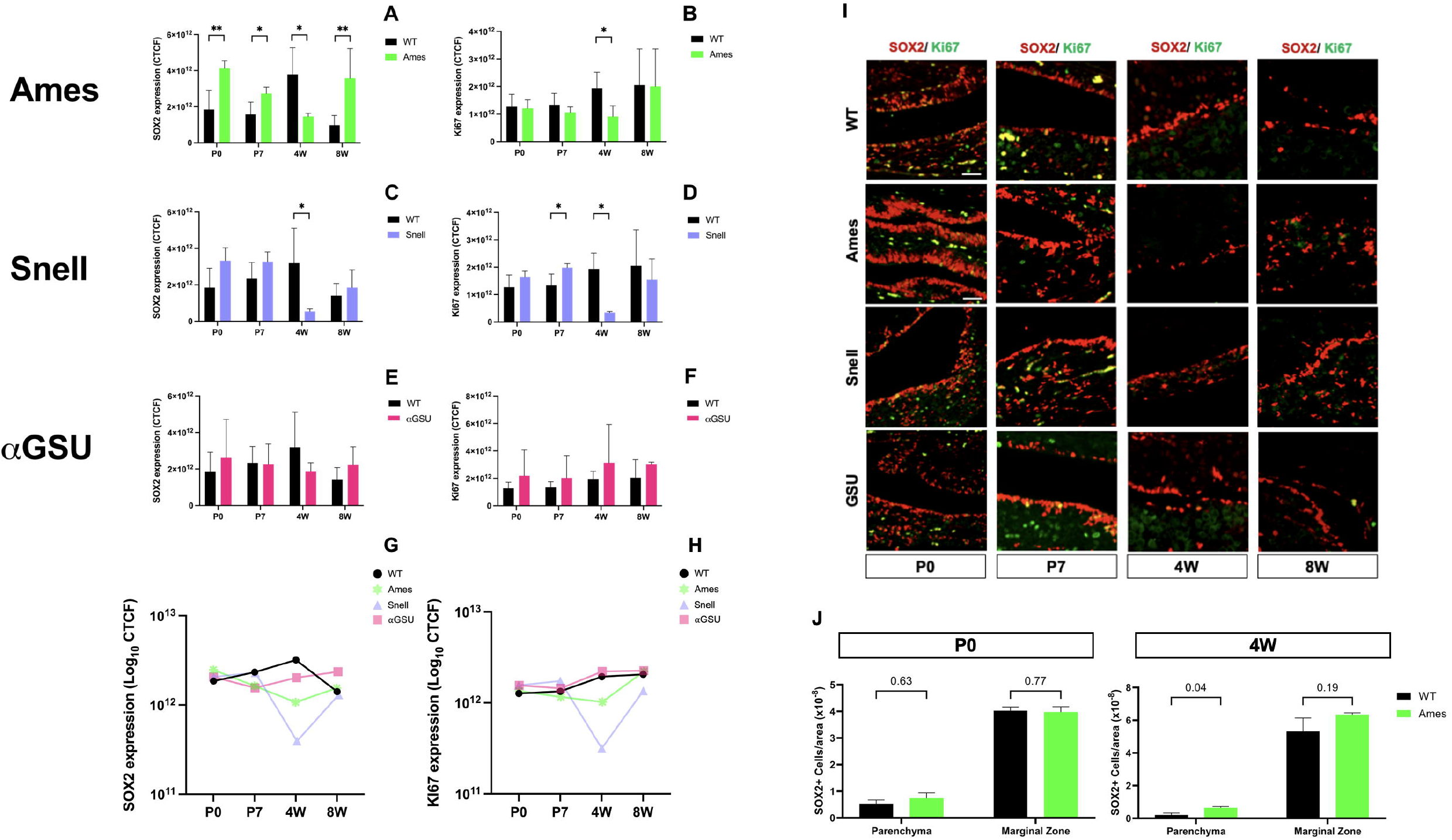
Elevated SOX2 protein in postnatal pituitaries of *Prop1-*deficient mice. Immunofluorescence intensity of SOX2 (A, C and E) and Ki67 (B, D and F) are expressed in log_10_ of corrected total cellular fluorescence (CTCF) for *Prop*^*df/df*^ (green), *Pou1f1*^*dw/dw*^ (purple), and αGSU mutant (pink) mice compared to WT (black) at postnatal periods. Results are expressed as the mean ± standard deviation (SD). Significant differences between groups are expressed as ^*^ p<0.05 and ^**^p<0.01. Timeline analysis of SOX2 and Ki67 expression in the wild-type (black), Ames (green), Snell (purple), and alpha-glycoprotein subunit (αGSU) (pink) mice (G and H) at postnatal periods. Results are expressed as the mean of each group. SOX2 (red) and Ki67 (green) immunofluorescence showing the marginal zone and parenchyma cell positivity (I). SOX2 positive cells proportion according to marginal zone or parenchyma localization in the WT (black) and Ames (green) mice at P0 and 4W (J).

The *Pou1f1* deficient mice had consistent levels of *Sox2* transcripts at all ages examined except at 8W, when there was an increase. There were no significant changes in *Ki67* transcripts. At the protein level, we confirmed that there were consistent levels of SOX2 protein at most ages, although there was a reduction at 4W (Figure 2C). The levels of KI67 protein were consistent between mutants and WT, except for a decrease at 4W (Figure 2D). It is noteworthy that both *Prop1* and *Pou1f1* deficient mice exhibit reduced SOX2 immunostaining at 4 wk relative to WT.

The αGSU deficient mice exhibited stable levels of *Sox2* and *Ki67* transcripts relative to wild type at all ages examined. We confirmed this at the protein level by demonstrating normal SOX2 and Ki67 protein levels at all ages (Figure 2E and F).

The temporal analysis shows similar SOX2 and Ki67 expression patterns in *Prop1* and *Pou1f1* mutants (and between the WT and αGSU models (Figure 2 G, H). For *Prop1* and *Pou1f1* mutants, these markers are significantly decreased at 4W compared to the other time points, while in WT and αGSU, SOX2 levels are stable over time with a slight decrease at 8W. SOX2 positive cells are mainly located in the marginal zone and Ki67 in the parenchyma of the gland (Figure 2I). The marginal zone of WT mice typically has a single layer of SOX2 positive cells, while the marginal zone of *Prop1* mutants has multiple SOX2 positive cells. This is expected because the stem cells fail to migrate out of the niche (reference Rob Ward’s paper). We quantified the number of SOX2 immunopositive cells in each region relative to the area and found no differences in *Prop1* mutants at P0 and 4W. This supports the idea that the stem cells are similar in size in mutants and WT.

### *Prop1* mutant pituitaries have differentiating gonadotrophs and lactotrophs that co-express SOX2

In order to understand the transcriptional changes observed in the *Prop1* mutant mice during the pubertal period (4W), we analyzed pituitary hormone expression and SOX2 co-localization, as well as hormone serum levels at two stages of sexual maturation. P30 represents the onset of sexual development, as assessed by vaginal opening and first vaginal cornification, in females, and these steps are completed by P40 (30). Normal cycling is typically established by 8W. For males, androgen-dependent preputial gland separation occurs P22-42, and by ∼P40 testosterone peaks, mature sperm are found in the testis, and mating behavior begins (31). We were able to detect GH, LHß, and PRL positive cells in the *Prop1* mutant pituitaries (Figure 3A, C and D), while none of the cells expressed TSHß (Figure 3B). Interestingly, a proportion of these hormone producing cell types co-expressed SOX2, suggesting that they might undergo differentiation, but it is not enough to properly secrete hormones and recover the phenotype (Figure 2E-H). Then, we accessed pituitary hormone serum levels at three different time points (P30, P40 and P60) to understand if the produced hormones are secreted in the bloodstream during the fourth week of *Prop1* mutant mouse development. At all periods, GH and FSH were significantly reduced in the serum of *Prop1* mutants (Figure 3I and K), while LH serum level dynamically changed during the sexual maturation period increasing at P40 (Figure 3J), suggesting that the pituitary tries to adapt, but cannot adequately support hormonal production and overcome the hypogonadism.

**Figure 3.**
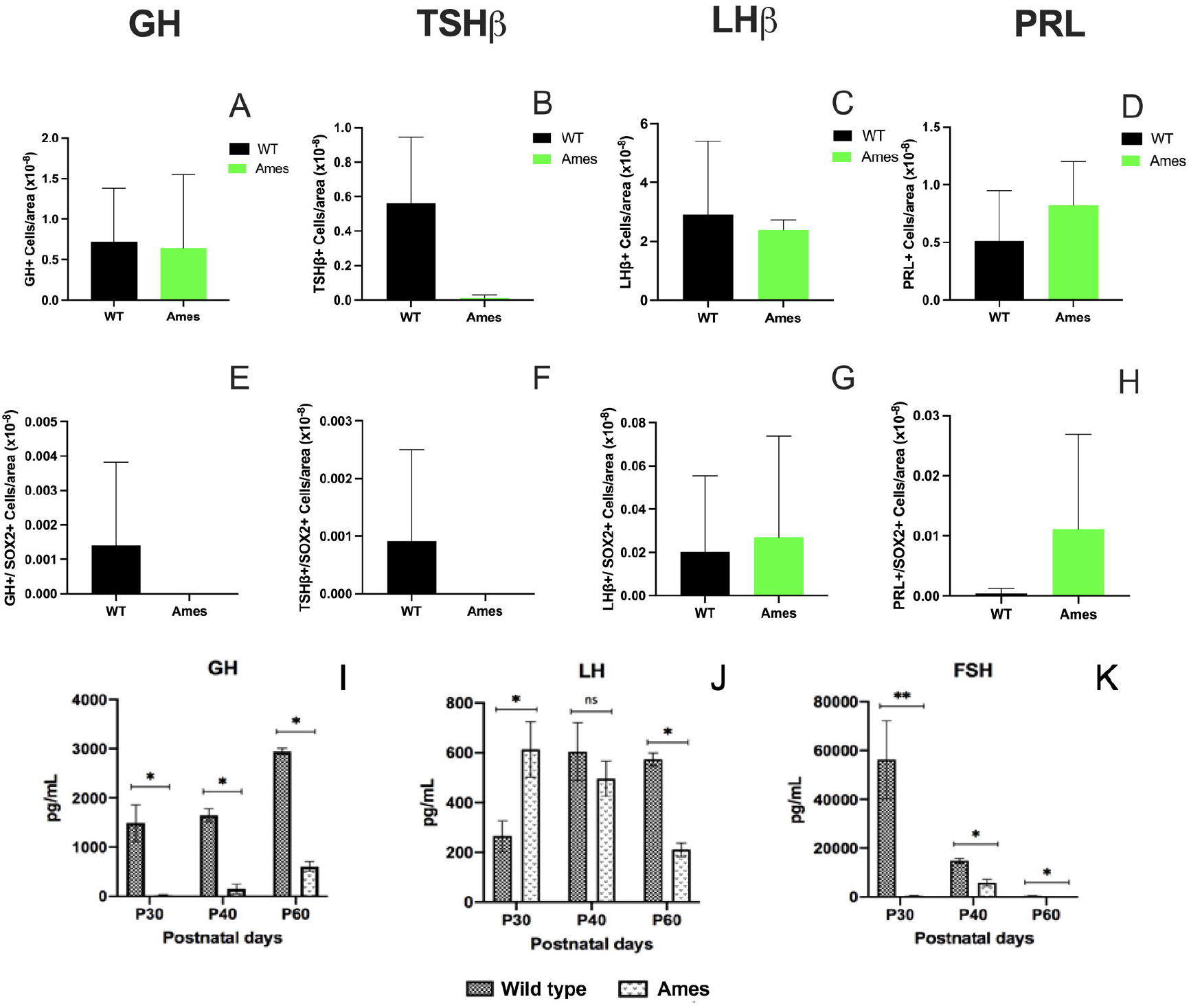
Colocalization of SOX2 and hormone proteins in *Prop1* mutants. Pituitary hormones GH, TSH β, LH β and PRL positive cells in WT (black) and *Prop1*^*df/df*^ (green) pituitaries (A-D). SOX2 and hormone co-localization in WT (black) and *Prop1*^*df/df*^ (green) pituitaries (E-H) at 4W. GH (I), LH (J) and FSH (K) serum levels in WT (dark) and *Prop1*^*df/df*^ (white) mice at P30, P40 and P60. Results are expressed as the mean ± standard deviation (SD). Significant differences between groups are expressed as ^*^ p<0.05.

## Discussion

Recent advances in multi-omics technologies have demonstrated the dynamic composition of pituitary cells and changes in pituitary gene expression regulating cell fate decisions and hormonal production (32-34). However, information regarding transcriptional dynamics of key pituitary development markers during postnatal pituitary development in animal models of hypopituitarism is lacking. We took advantage of the three previously reported mouse strains (*Prop1*^*df/df*^, *Pou1f1*^*dw/dw*^ and αGSU mutant) with hypopituitarism harboring distinct molecular defects and aimed to assess the effect of these genetic variants on the expression of pituitary stem cell markers, transcription factors, and cell proliferation markers expression at key timepoints of postnatal development. We also performed SOX2 immunofluorescence to correlate the mRNA and protein changes in the specific pituitary stem cell markers during the postnatal pituitary development. After analyzing the transcriptional data, we decided to take a closer look at the most interesting differences found, which were in the pubertal period of the *Prop1*^*df/df*^ mice. We identified changes in SOX2 expression that were divergent from the ones in *Pou1f1*^*dw/dw*^ and αGSU mice. The onset of puberty in mice is expected to occur between P28 and P40 (31). Looking at early and late timepoints of sexual maturation, we identified a dynamic trend in pituitary stem cell marker and gonadotropin expression in the *Prop1*^*df/df*^ model of hypopituitarism showing that, even with no restoration of hypogonadal phenotype, this animal can produce and secrete hormones.

Pituitary stem cell markers, *Sox2* and *Sox9*, showed progressively increased expression during postnatal development in *Prop1*^*df/df*^ and *Pou1f1*^*dw/dw*^ mice (Figure1, S2 and S4). Since elevated *Sox2* expression could be due to fewer differentiated cells, SOX2 immunostaining in mutant and wild-type pituitaries was quantified at the four postnatal periods to determine SOX2-positive cells. One possible explanation for the generally increased expression of the stem cells markers in *Prop1* mutant mice is the failure to progress to activate *Pou1f1* expression. At most ages *Sox2* expression is similar in normal and *Pou1f1*^*dw/dw*^ pituitaries (16). While both *Pou1f1* and *Prop1* are essential for pituitary organogenesis and cell specification, *Prop1* is co-expressed in pituitary SOX2-positive stem cells and is necessary for cell cycle regulation and cell migration; hence, the absence of these pituitary transcription factors could keep cells in an undifferentiated state (3, 16, 19).

*Hesx1* encodes a transcription factor whose expression is regulated by *Lhx3* and *Prop1* (35, 36). *Hesx1* is co-expressed with *Lhx3* at the initial stage of Rathke’s pouch development, primarily in stem cells, and it is down regulated prior to differentiation of hormone-producing cells (28). Increased *Hesx1* expression in the *Prop1* mutant was expected because *Prop1* and β-catenin (CTNNB1) are necessary for repression of *Hesx1* expression (28, 37). Since *Prop1* represses *Hesx1*, and activates *Pou1f1*, no changes in *Hesx1* expression were expected in *Pou1f1* mutants. Indeed, *Hesx1* expression in *Pou1f1* mutants was similar to wild type at most ages (P0-4wk).

*Prop1*, but not *Pou1f1*, mutant pituitaries present significant dysmorphology during embryogenesis due to the lack of stem cell migration out of the niche in *Prop1* mutants (38, 39). Both *Prop1* and *Pou1f1* mutants have hypoplastic pituitaries by P8 and P11, respectively, which has been attributed, in part, to reduced cell proliferation (38). We confirmed reduced *Ki67* mRNA expression at P7 and 4W, respectively.

In the αGSU mice, there was no difference in *Sox2* expression in mutant and wild-type animals at any of the ages examined, suggesting no effect of this mutation on the stem cell pool. Similarly, there was no difference in cell proliferation markers or transcription factors. This is consistent with the ability of these mutant stem cells to differentiate into all pituitary hormone producing cell types when thyroid hormone is replaced. The hypopituitarism in these mutants arises because none of the heterodimeric glycoprotein hormones (TSH, LH and FSH) can be secreted without the alpha subunit (40).

At the analyzed periods of sexual development, we also observed a dynamic in SOX2 expression and gonadotropins, which has never been previously reported. SOX2 plays an important role in pituitary development and is the main marker of pituitary stem cells (3, 4). Pituitary can adapt its cell composition during physiological demand and injury, including the pubertal period (5-8). Despite the unexpected increase in gonadotropin mRNAs and proteins during puberty of *Prop1* mutants, fertility is not restored in our mutant colony. Some studies have demonstrated occasional cases of fertile male *Prop1* mutant mice (41), which is not fully understood or explained. It is possible that the penetrance of male fertility in mutants is influenced by genetic background and/or high fat diet.

In conclusion, this study is the first to analyze mRNA expression levels of the pituitary stem cell markers, proliferation markers, and pituitary transcription factors during four periods of postnatal development in established mouse models of congenital hypopituitarism. The transcriptional expression dynamics of the stem/progenitor pituitary cell markers, cell proliferation markers, and pituitary transcription factors change depending on the molecular defect. Alterations in the transcription factors expressed early in the differentiating pituitary (*Prop1* and *Pou1f1*) promote an increased expression of the pituitary stem cell marker *Sox2*. However, molecular defects in the hormone subunit gene *Cga* do not affect the expression levels of these cell markers. Thus, our data facilitate the understanding of key pituitary gene transcriptional dynamics in different molecular defects and periods. Finally, this study is showed transient gonadotropin production and secretion during *Prop1* mice sexual maturation, even though phenotype was not recovered.

## Supporting information

Assessment of postnatal transcription of Hes1, Notch2, and Otx2 in hypopituitary mutants postnatally.

Temporal changes in Hes1, Notch2 and Otx2 expression in hypopituitary mutants

RT-qPCR raw data

## Abbreviations

SOX2: SRY-box transcription factor 2
SOX9: SRY-box transcription factor 9
SOX3: SRY-box transcription factor 3
S100β: S100 protein, beta polypeptide, neural
HES1: Hairy and enhancer of split-1
TLE: Transducin-like enhancer of split
HESX1: Homeobox gene in embryonic stem cell 1
OTX2: Orthodenticle homeobox 2
LHX3: LIM homeobox 3
PROP1: PROP paired-like homeobox 1
POU1F1: POU class 1 homeobox 1
NOTCH2: Notch receptor 2
CGA: Glycoprotein hormones, alpha polypeptide
Ki67: Marker of proliferation Ki-67
CTNNB1: Catenin beta 1
TSH: Thyroid-stimulating hormone
LH: Luteinizing hormone
FSH: Follicle-stimulating hormone
GH: Growth hormone
ACTH: Adrenocorticotropic hormone
PRL: Prolactin

## Acknowledgments

This project was possible for the donation of three strains of animals. by Prof. Dr. Sally A. Camper at the University of Michigan.

## Authorship confirmation statement

Juliana Moreira Silva: Investigation, methods, formal analysis, writing—original draft

Claudia V Chang: Conceptualization, investigation

Ricardo V Araujo: Investigation, methods

Cinthya S Cerqueira: Methods

Nicholas Trigueiro: Methods, statistical analysis

Bruna Viscardi de Azevedo: Investigation, methods

Berenice Bilharinho de Mendonça: Funding acquisition, supervision

Nicholas Hoch: Methods and review

Lilian Cristina Russo: Methods Luciani R S Carvalho: Conceptualization, funding acquisition, project administration, supervision, writing—review and editing.

## Figure legends

**S1**. <u>Assessment of postnatal transcription of</u>*<u>Hes1,Notch2</u>*<u>, and</u>*<u>Otx2</u>*<u>in hypopituitary mutants postnatally.</u>RT-qPCR results expressed in log_2_ ^Δ^^CT^ of the genes *Hes1* (A, D and G), *Notch2* (B, E and H) and *Otx2* (C, F and I) in the wild-type, *Prop1*^*df/df*^, *Pou1f1*^*dw/dw*^, and alpha-glycoprotein subunit (αGSU) mice at postnatal periods (newborn: P0, one week: P7, four weeks: 4W, and eight weeks: 8W). Results are expressed as the mean ± standard deviation (SD). Asterisks indicate significant differences between groups (p < 0.05).

**S2**. <u>Assessment of postnatal transcription of</u>*<u>S100β, Sox3</u>*<u>, and</u>*<u>Sox9</u>*<u>in hypopituitary mutants postnatally.</u>RT-qPCR results expressed in log_2_ ^Δ^^CT^ of the genes *S100* β (A, D and G), *Sox3* (B, E and H) and *Sox9* (C, F and I) in the wild-type, *Prop1*^*df/df*^, *Pou1f1*^*dw/dw*^, and alpha-glycoprotein subunit (αGSU) mice at postnatal periods (newborn: P0, one week: P7, four weeks: 4W, and eight weeks: 8W). Results are expressed as the mean ± standard deviation (SD). Asterisks indicate significant differences between groups (p < 0.05).

**S3**. Temporal changes in *Hes1, Notch2* and *Otx2* expression in hypopituitary mutants. Timeline of *Hes1, Notch2* and *Otx2* expression in the wild-type (black), *Prop1*^*df/df*^ (green), *Pou1f1*^*dw/dw*^ (purple), and alpha-glycoprotein subunit (αGSU) (pink) mice (J to L) at postnatal periods. Results are expressed as the mean of each group.

**S4**. Temporal changes in *<u>S100β, Sox3</u>*<u>, and</u>*<u>Sox9</u>* expression in hypopituitary mutants Timeline of *S100* β, *Sox3* and *Sox9* expression in the wild-type (black), *Prop1*^*df/df*^ (green), *Pou1f1*^*dw/dw*^ (purple), and alpha-glycoprotein subunit (αGSU) (pink) mice (J to L) at postnatal periods. Results are expressed as the mean of each group.

**S5**. RT-qPCR raw data

