## Supplementary figures and images for "Temporal Analysis of Pituitary Transcriptional Dynamics in Mice Models of Hypopituitarism During Postnatal Development"

### Assessment of postnatal transcription of Hes1, Notch2, and Otx2 in hypopituitary mutants postnatally.

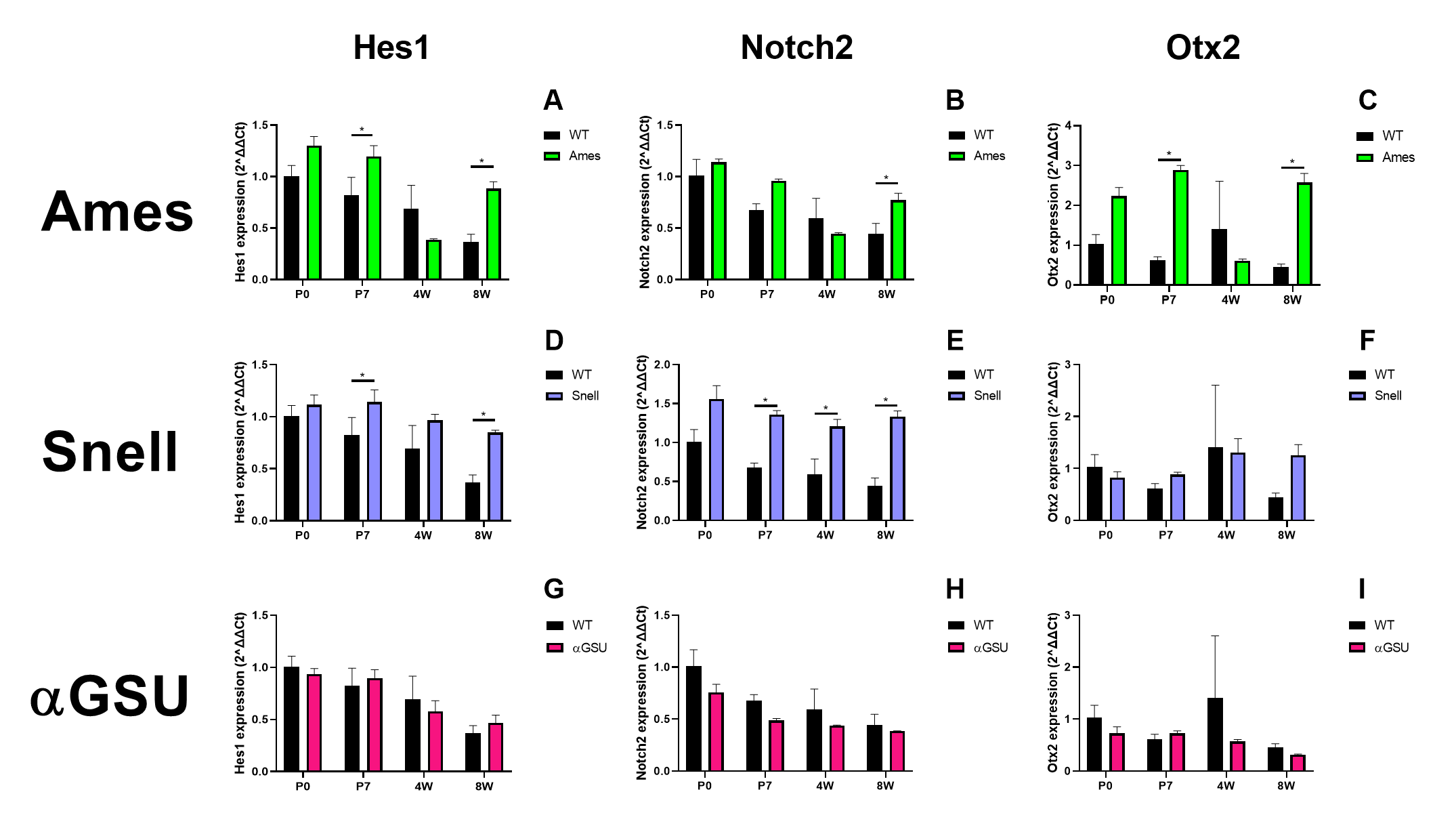

### Supplemental Data 1

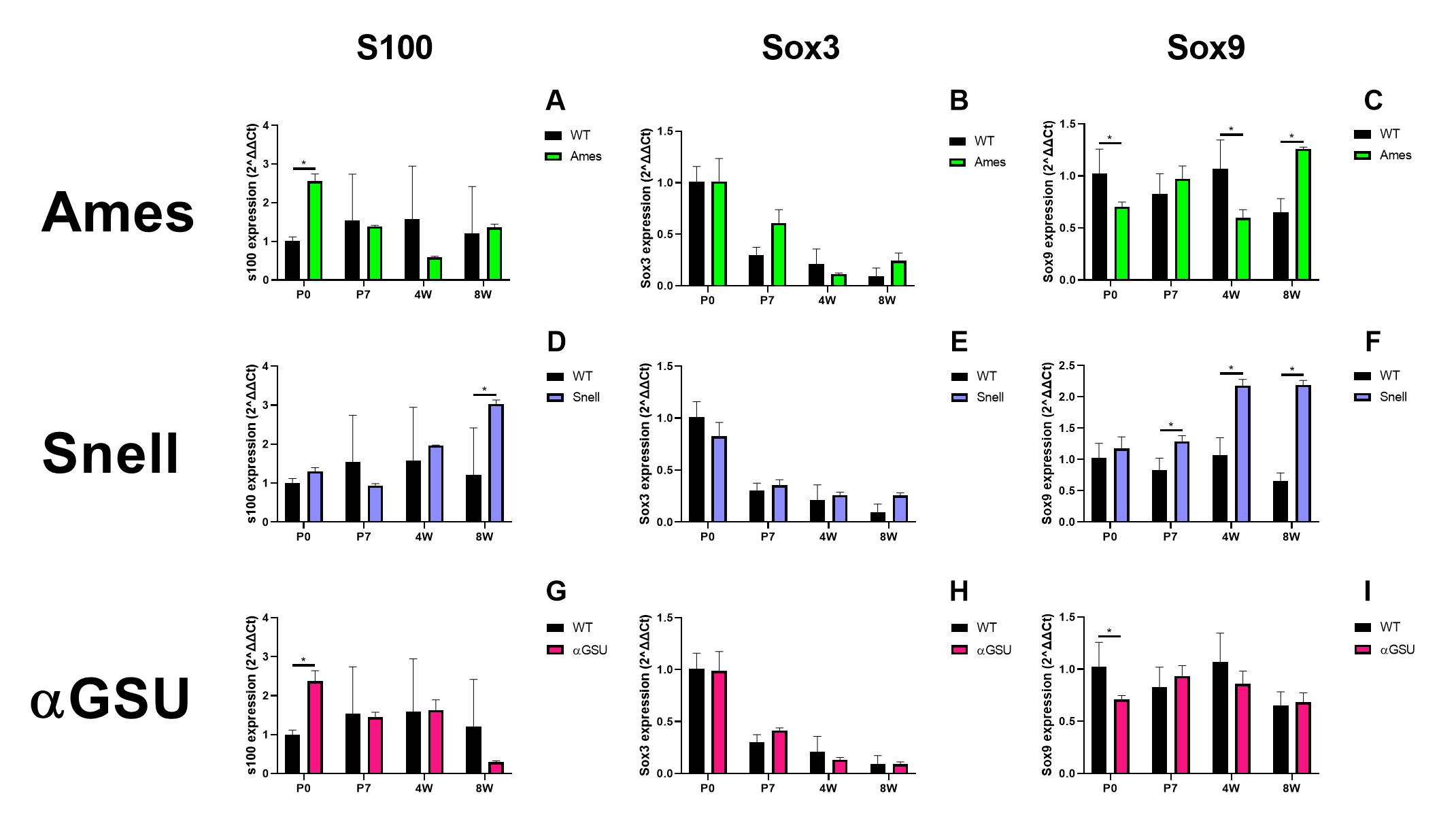

### Supplemental Data 2

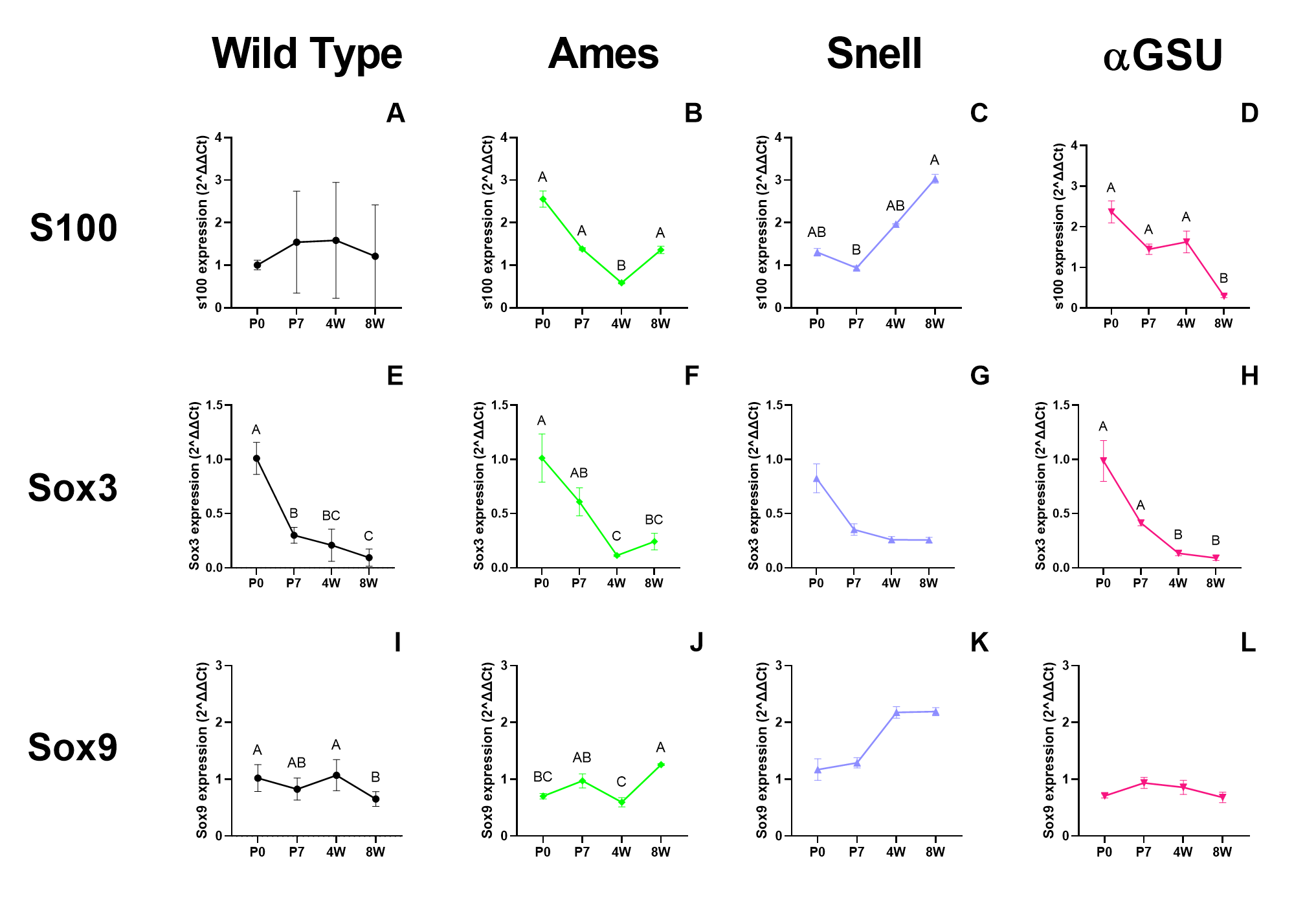

### Temporal changes in Hes1, Notch2 and Otx2 expression in hypopituitary mutants

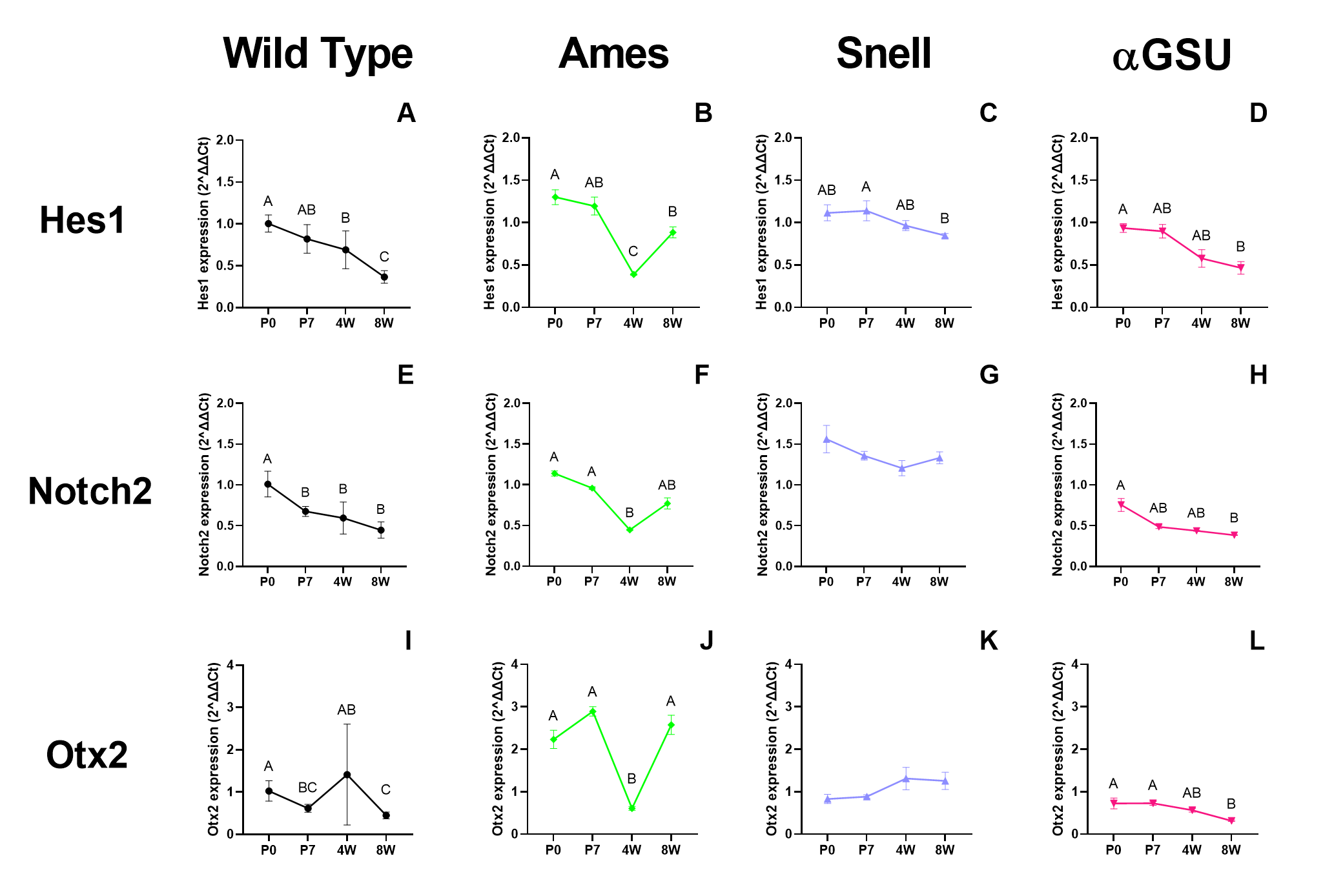
